# Five days of heat acclimation improves cardiovascular and thermoregulatory responses without altering renal stress biomarkers in endurance athletes

**DOI:** 10.64898/2026.03.06.710014

**Authors:** Daniel Snape, Barney Wainwright, Iain T Parsons, Michael J Stacey, David R Woods, John O’Hara

**Author notes:** **Corresponding author** Daniel Snape. **Statements and declarations**. **Data Availability** The dataset analysed during the current study may be requested upon an informal inquiry addressed to the corresponding author. **Competing Interests** The authors have no relevant financial or non-financial interests to disclose. **Author Contribution Statement** DS conceived the study, collected results, performed data analysis, and drafted the manuscript. IP collected results and reviewed and revised the manuscript. MS and DW critically edited the manuscript and contributed to the content. BW and JOH contributed to the concept and design, review and revised the manuscript, and contributed to data interpretation.

## Abstract

Short-term heat acclimation (HA) induces cardiovascular and fluid-regulatory adaptations, but it’s impact on markers of renal tubular injury and acute kidney injury risk (AKI) during exercise-heat stress remains unclear. Fourteen healthy endurance athletes were randomised to five days of isothermic HA (HOT; *n* = 7; 32 °C, 70% relative humidity; target core temperature ≥38.5 °C), or matched exercise in thermoneutral conditions (TEMP, *n* = 7). Heat stress tests (HST; 45 min cycling at 32 °C, 70% RH) were performed pre- and post-intervention. Blood biomarkers of kidney tubular stress (NGAL, KIM-1), fluid-regulation (copeptin, serum osmolality) and sympathetic activity (plasma normetanephrine) were measured at rest and immediately post-HST. HA reduced resting heart rate (-8 ± 5 bpm, *p* = 0.007, *d* = 1.0), increased plasma volume (+7.3 ± 5.1%, *p* = 0.022) and sweat loss (+500 ± 539 mL, *p* = 0.018, *d* = 1.1). Copeptin rose during the pre-intervention HST in both groups (HOT: +11 ± 6; TEMP: +12 ± 13 pmol·L^−1^, *p* < 0.05), but not post-intervention. NGAL increased only in TEMP during HST1 (+45 ± 29 μg·L^−1^, *p* = 0.030), while KIM-1 remained unchanged. No group x time interactions were observed for any biomarkers (p > 0.05). Five days of HA improved cardiovascular and thermoregulatory responses but did not alter renal stress markers or fluid-regulatory responses during exercise in the heat. These findings suggest short-term HA enhances heat tolerance without reducing acute renal biomarker responses under hot, humid conditions.

**New & Noteworthy:** Five days of isothermic heat acclimation improved cardiovascular and thermoregulatory responses, related to a lower resting heart rate, plasma volume expansion, and greater sweat loss. However, these benefits did not reduce renal tubular stress markers (NGAL, KIM-1), fluid-regulatory strain (copeptin), or sympathetic activity (normetanephrine) during exercise in the heat. Short-term heat acclimation lowers cardiovascular strain but does not mitigate renal biomarker responses, suggesting kidney stress risk remains unchanged in hot, humid conditions.

## Introduction

Endurance athletes frequently train and compete in hot, humid environments where metabolic heat production and environmental heat stress impose significant thermal and cardiovascular strain. Inadequate heat dissipation elevates core temperature, impairing performance and increasing risk of heat-related illness, including exertional heat stroke, which can lead to systemic inflammation and multi-organ dysfunction (Leon & Bouchama, 2015). During exercise, redistribution of cardiac output to active muscle and skin reduces renal blood flow by 25–50% compared with rest, an effect exacerbated in the heat as skin blood flow and sweating rates increase (Schlader et al., 2019). Fluid loss through sweat and plasma volume contraction further diminishes renal perfusion, while neurohumoral activation via sympathetic outflow, catecholamine release, and arginine vasopressin (AVP) secretion acts to conserve fluid but simultaneously exacerbates renal vasoconstriction (Patterson et al., 2004b). These responses increase tubular strain, induce transient renal dysfunction, reflected by elevated serum creatinine and reduced glomerular filtration rate consistent with AKI (Chapman et al., 2021; Pryor et al., 2020; Roncal-Jimenez et al., 2016).

Significant rises in serum creatinine have been reported following endurance events, often exceeding the diagnostic threshold for stage I AKI after marathons, ultra-endurance races, and long-distance cycling (Lippi et al., 2011; Mansour et al., 2017). Creatinine often rises 24-36 hours after injury and can be confounded by exercise due to haemoconcentration and muscle metabolism rather than kidney injury, complicating interpretation in athletic cohorts (Liu et al., 2025). Therefore, collection of sensitive tubular stress biomarkers such as neutrophil gelatinase-associated lipocalin (NGAL) and kidney injury molecule-1 (KIM-1) during exercise-heat stress, which rise before changes in creatinine, may offer a more responsive insight into renal strain (Chapman et al., 2021; Pryor et al., 2020). In addition, collection of copeptin (a stable surrogate for AVP), plasma normetanephrine (reflecting sympathetic tone), and serum osmolality (a marker of hydration and osmotic stress) alongside NGAL and KIM-1 may provide an integrated view of systemic and renal responses to heat stress (Katan et al., 2008; Stacey et al., 2018; Stacey et al., 2018).

Improved plasma volume and sympathetic modulation following HA may preserve renal blood flow, reducing renal strain during exercise in the heat. These cardiovascular adaptations could offset the renal hypoperfusion and neurohumoral vasoconstriction that occur during exercise-heat stress. However, whether short-term HA confers such renal protection remains unclear. Controlled isothermic HA, which maintains core temperature ≥38.5 °C for 60-90 min per session, provides a potent and time-efficient stimulus for athletes and military personnel (Gibson et al., 2015). Five consecutive isothermic sessions are sufficient to lower core body temperature, expand plasma volume, improve sweating efficiency, and lower exercising heart rate (Tyler et al., 2024), with diminishing marginal returns thereafter (Moss et al., 2020). While these adaptations enhance thermal tolerance, there influence on renal perfusion and tubular stress remains unclear. Recent work by Connor et al., (2025) demonstrated that plasma NGAL and creatinine appear to be unaffected by acute heat stress before and after 10-days of fixed-intensity treadmill walking in a climatic chamber (40 °C) to induce HA. Although HA didn’t attenuate the rise in creatinine or NGAL, the kidneys exhibited an increased sensitivity to hyperthermia in heat-acclimated individuals.

Therefore, this study investigated whether a short-term (5-day, 90 min·day^−1^) isothermic HA protocol in endurance-trained individuals influences thermoregulatory, cardiovascular, renal, and neuroendocrine responses to exercise in the heat. We hypothesised that HA would reduce thermal and cardiovascular strain and attenuate biomarkers of renal tubular injury (NGAL, KIM-1), sympathetic activation (normetanephrine), and fluid-regulatory stress (copeptin and osmolality).

## Methods

### Ethics statement and participant population

The study protocol gained institutional ethical approval (Reference: 64628, Leeds Beckett University) and was conducted in line with the principles expressed in the *Declaration of Helsinki*, 2013. Fourteen endurance-trained athletes volunteered for the study (table 1). All participants were healthy, had not been exposed to hot conditions (>25 °C) for a least 3-months, and provided written informed consent. All individuals completed a minimum of 5h aerobic exercise per week and according to classifications by De Pauw et al., (2013) met the criteria for performance levels 2 or 3, see table 1.

**Table 1.** Baseline physical characteristics of the heat acclimation (HOT) and temperate exercise intervention (TEMP).

| Variable | HOT (*n* = 7) | TEMP (*n* = 7) | p-value |
| --- | --- | --- | --- |
| Age (years) | 29 ± 10 | 28 ± 9 | 0.780 |
| Height (cm) | 180 ± 5 | 179 ± 5 | 0.690 |
| Body mass (kg) | 73.4 ± 6.5 | 75.6 ± 10.0 | 0.646 |
| VO<sub>2max</sub> (mL.min.kg<sup>-1</sup>) | 63 ± 8 | 55 ± 6 | 0.051 |
| MRMP (watts) | 400 ± 37 | 339 ± 36 | 0.011* |

*Note:* Values represent the mean ± SD. The reported values of maximal oxygen consumption (VO<sub>2max</sub>), maximum ramp minute power (MRMP), were measured under temperate conditions (20 °C, 45% relative humidity). \*Significant difference (p = 0.011) between groups in MRMP.

### Design of study

A randomised, between-groups, pre-post design was employed. Participants completed five consecutive daily intervention sessions (HOT or TEMP). Heat stress tests (HSTs) were conducted exactly 48 hours before the first intervention session (HST1) and 48 hours after the final session (HST2). Randomisation to groups was performed using GraphPad Prism software. Testing occurred between 07:00-11:00am to control for circadian variation and participants arrived euhydrated (urine osmolality <700 mOsmol kg^−1^; urine specific gravity <1.020). Training temperatures were specifically chosen to mimic the conditions athletes were anticipated to face during the Tokyo 2020 games (30-35 °C and 70-90% RH) (Kakamu et al., 2017).

### Preliminary testing

Participants’ stature (Seca, 220, Hamburg, Germany) and body mass (Seca, 770) were initially measured. A graded exercise test (GXT) was performed on a cycle ergometer (Wattbike, Trainer, Nottingham, UK) under temperate laboratory conditions (19.3 ± 1.6 °C, 46 ± 6% RH). Participants cycled at 150 W, increasing by 25 watts (W) every 4 min until blood lactate concentration exceeded 4.0 mmol L^−1^, measured on a biochemistry analyser (EKF, Biosen C-Line, Cardiff, UK). Following a 15-min recovery, participants cycled at 150 W and increased the power by 20 W min^−1^ until volitional exhaustion. Heart rate (HR) was recorded continuously via chest strap telemetry (V800, Polar, Kempele, Finland) and expired gases were measured using a calibrated online gas analysis system (Cortex, Metalyzer 3B, Leipzig, Germany). Maximal oxygen uptake (VO_2max_) was determined according to standard criteria of Winter et al., (2006).

### Heat stress test

HST’s involved 45 minutes cycling at 2.5 W.kg^-1^ within an environmental chamber (Sporting Edge, UK) set to hot-humid conditions (32.0 °C ± 0.2, 70.4 ± 0.8% RH), adapted from (Snape et al., 2023). Gastrointestinal temperature (Tgi) was measured via a telemetric pill (e-Celsius, BodyCap, Caen, France), calibrated through water immersion. Skin temperature (Tskin) was measured via four iButton sensors (Maxim Integrated Products, DS1922L, CA, USA), and mean Tskin calculated using the equation by Ramanathan, (1964). Sweat patches (3M, Tagaderm +Pad, Minnesota, USA) were applied to the forearm for the collection of localised sweat. During exercise HR was monitored continuously, Tre at 2.5-min intervals, and perceived exertion (RPE; Borg, (1982)), thermal sensation (TSS; Toner et al., (1986)) and thermal comfort (TC; Zhang et al., (2004)) recorded at 5-min intervals. Sweat loss (mL) was estimated from nude body mass changes, corrected for urine output. Venous blood samples were collected pre-and post-HST for analysis of NGAL, KIM-1, copeptin, normetanephrine, and serum osmolality. Capillary blood samples were analysed in triplicate for haemoglobin (Hb) and haematocrit (Hct) to calculate plasma volume (PV) changes.

### Heat acclimation intervention (HOT)

Participants in the HOT group completed five consecutive 90-minute sessions in a hot-humid environment (32.2 ± 0.2 °C, 69.7 ± 1.8% RH) using the controlled hyperthermia methodology (Fox et al., 1967). Exercise intensity was set at 2.5 W.kg^-1^ to raise core temperature to 38.5 °C within ∼30-minutes, followed by passive or reduced-intensity exercise to maintain target temperature. Heart rate and perceptual measures (RPE, TSS, TC) were recorded throughout. Fluid intake was monitored, with bottle weight measured pre-to post-session to the nearest 0.1 g using weighing scales (Ohaus, C series, New Jersey, USA).

### Temperate climate exercise intervention (TEMP)

TEMP participants completed five sessions in a thermo-neutral environment (21.3 ± 1.3 °C, 37.3 ± 6.1% RH) at a matched workload. Physiological and perceptual measures were recorded at similar intervals. Fluid intake was *ad libitum* and recorded to calculate sweat loss.

### Analytical methods

Changes in plasma volume (ΔPV) were calculated using Dill & Costill, (1974) equation from measured Hct and Hb. Sweat Na^+^ concentration was measured using a flame photometer (Jenway, PFP7 Flame Photometer, IL, USA). Venous blood samples were processed in serum separator (Becton Dickinson, SSTTM II Advance, Franklin Lakes, NJ, USA) and EDTA tubes (32.332 Sarstedt, Akteingesellscaft & Co., Numbrecht, Germany), centrifuged at 2600 rpm for 15-minutes at 4 °C, and aliquots stored at -80 °C. Serum osmolality was measured in duplicate by a micro-osmometer (Advanced Model 3320, Advanced Instruments, USA; CV 1.1%). Normetanephrine was analysed via LC-MS/MS (CV: 4–12%), copeptin via immunoassay (Brahms, CT-proAVP Kryptor Compact Plus, USA; CV: 2–10%). KIM-1 and NGAL were analysed using commercially available ELISA kits (R & D systems Europe, Abingdon, Oxon) by a specialist biomarker laboratory (Affinity Biomarker Labs, London). Intra/inter assay variability for KIM-1 was (CV: 1.3-7.0%), and for NGAL (CV: 2.3-3.9%). The lower limit of detection was 1.31 ng. L^-1^, and 0.012 µg/L^-1^, respectively. Hormone concentrations presented are uncorrected for changes in plasma volume.

### Statistical Analysis

Prior to experimentation, an *a-priori* power calculation was performed using G*Power software (version 3.1) to determine the sample size required. Using previously reported data where the primary outcome variables were the same as the present study (i.e., gastrointestinal temperature, whole body sweat loss, heart rate), the minimum effect size reported to observe a difference was 0.50. A total number of 12 participants was deemed as being necessary based upon an alpha (α: 0.05) and beta (β: 0.80).

Statistical calculations were performed using GraphPad Prism (version 9.0). Data are reported as means ± standard deviation (SD) and significance set at *P <* 0.05. Normality was assessed using the Shapiro–Wilk test (*P* ≥ 0.05). Parametric data from the pre-intervention HST were analysed using an independent-samples *t-test* between groups (HOT and TEMP), while non-parametric perceptual data were analysed using a Wilcoxon signed-rank test. The effects of time (PRE or POST) and group (HOT or TEMP) on continuous variables were determined using a two-way ANOVA for repeated measures. Where data were non-parametric but log-transformed data were parametric, ANOVA was performed on the log values. If log transformation did not normalize data, ANOVA was applied only when the Browne-Forsythe test indicated equal variance. Significant interactions were followed by Bonferroni-corrected post-hoc tests (alpha = 0.05). Effect sizes (Cohen’s d) were calculated to assess the magnitude of differences (Lakens, 2013), categorized as trivial (<0.2), small (>0.2), moderate (>0.6), large (>1.2), very large (>2.0), and extremely large (>4.0).

## Results

Fourteen endurance-trained participants were randomised to HOT (*n* = 7) or the TEMP (*n* = 7). All completed the five-day intervention and both heat stress tests (HST1 and HST2), except one participant in the TEMP did not complete venous blood sampling due to trypanophobia.

### Heat stress testing – physiological and perceptual adaptations

Baseline hydration indices did not differ between groups (urine osmolality: *p* = 0. 839, *d* = 0.1; urine specific gravity: *p* = 0.352, *d* = 0.6; serum osmolality: p = 0.369, *d* = 0.6). A summary of resting and peak physiological measures throughout each intervention are presented in table 2. Resting HR showed a significant group × time interaction (P = 0.014) and main effect of time (P = 0.047). Post-hoc analysis revealed a moderate reduction in HOT (-8 ± 5 bpm, P = 0.007, d = 1.0), with no change in TEMP. Resting Tgi and Tskin were unchanged in both groups (P > 0.05). Plasma volume increased significantly in HOT (+7.3 ± 5.1%, P = 0.022) but not TEMP (+3.0 ± 6.4%, P = 0.393). Sweat loss demonstrated a significant interaction (P = 0.010), with HOT showing a moderate increase (+500 ± 539 mL, P = 0.017, d = 1.1). No interaction was observed for peak RPE, TSS, or TC (P > 0.05). However, time effects indicated improvements in HOT for peak RPE (-2 ± 2, P = 0.007, d = 1.0) and TC (-1 ± 1, P = 0.016, d = 1.3). TEMP showed no significant changes.

**Table 2.** Physiological and perceptual responses at rest and during cycling heat stress tests prior to either heat acclimation (HOT) or temperature exercise intervention (TEMP).

|  |  | HOT (*n* = 7) |  |  | TEMP (*n* = 7) |  |  | Interaction |
| --- | --- | --- | --- | --- | --- | --- | --- | --- |
|  |  | HST1 | HST2 | Δ HST | HST1 | HST2 | Δ HST | *p*-value |
| Heart rate (bpm) | REST | 63 ± 6 | 55 ± 9 | -8 ± 5* | 61 ± 5 | 62 ± 7 | +1 ± 6 | 0.014 |
|  | PEAK | 168 ± 9 | 163 ± 10 | -5 ± 3 | 176 ± 12 | 175 ± 11 | -1 ± 6 | 0.264 |
| T<sub>gi</sub> (°C) | REST | 36.7 ± 0.2 | 36.5 ± 0.2 | -0.2 ± 0.3 | 36.7 ± 0.3 | 36.6 ± 0.3 | -0.1 ± 0.1 | 0.586 |
|  | PEAK | 38.9 ± 0.6 | 38.7 ± 0.4 | -0.2 ± 0.3 | 38.6 ± 0.6 | 38.6 ± 0.6 | 0 ± 0.3 | 0.687 |
| T<sub>skin</sub> (°C) | REST | 33.7 ± 0.5 | 33.5 ± 0.3 | -0.2 ± 0.3 | 33.8 ± 0.6 | 33.7 ± 0.7 | -0.1 ± 0.2 | 0.934 |
|  | PEAK | 36.2 ± 0.2 | 36.1 ± 0.3 | -0.1 ± 0.4 | 36.7 ± 0.3 | 36.8 ± 0.3 | 0.1 ± 0.1 | 0.065 |
| RPE (AU) | PEAK | 16 ± 2 | 14 ± 2 | -2 ± 2 | 18 ± 2 | 17 ± 2 | -1 ± 1 | 0.257 |
| Thermal sensation (AU) |  |  |  |  |  |  |  |  |

### Renal and neuroendocrine blood biomarkers following intervention

There were no significant group × time interactions for any blood biomarker (P > 0.05), see figure 1. Post-HST concentrations of NMET (-5.7%, P > 0.999, d = 0.2), copeptin (-21.5%, P = 0.314, d = 0.5), NGAL (-13.9%, P = 0.716, d = 0.7), and KIM-1 (+22.3%, P > 0.808, d = 0.2) were unchanged after HA. TEMP also showed no changes (P > 0.05).

**Figure 1.**
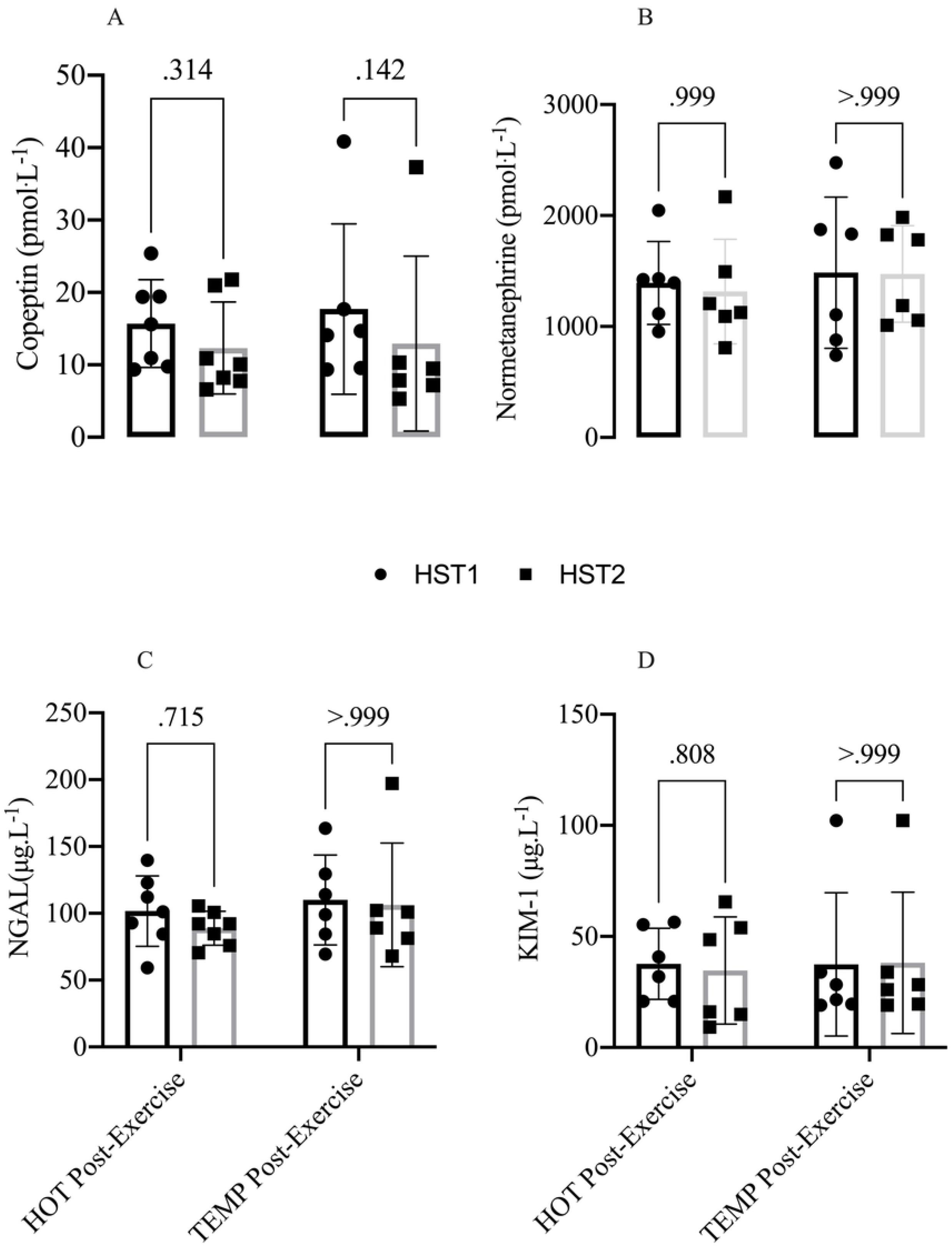
Mean (SD) and individual data points for (A) copeptin, (B) normetanephrine, (C) neutrophil-gelatinase associated lipocalin (NGAL), and (D) kidney injury molecule-1 (KIM-1) concentrations measured immediately post-exercise during heat stress tests (HST1: pre-intervention; HST2: post-intervention) in the heat acclimation group (HOT) and temperate climate exercise group (TEMP). Two-way ANOVA results for copeptin: Group (p = 0.794), Time (p = 0.030), Interaction (p = 0.672); NMET: Group (p = 0.664), Time (p = 0.585), Interaction (p = 0.679); NGAL: Group (p = 0.383), Time (p = 0.422), Interaction (p = 0.646); KIM-1: Group (p = 0.919), Time (p = 0.652), Interaction (p = 0.461). Pairwise comparisons shown above bars.

### Acute response of blood biomarkers to exercise-heat stress

Plasma NMET increased significantly from PRE to POST in both groups during HST1 (HOT: +990 ± 1025 pmol·L^−1^; TEMP: +1106 ± 1156 pmol·L^−1^; P < 0.001) and HST2 (HOT: +918 ± 965 pmol·L^−1^; TEMP: +1035 ± 1062 pmol·L^−1^; P < 0.001). Copeptin rose during HST1 in both groups (HOT: +11 ± 6 pmol·L^−1^, P = 0.015; TEMP: +12 ± 13 pmol·L^−1^, P = 0.019) but not HST2 (P > 0.05). Serum osmolality was similar between groups at rest (HST1: HOT 295 ± 6 vs TEMP 293 ± 4 mOsm·kg^−1^; HST2: HOT 291 ± 4 vs TEMP 295 ± 5 mOsm·kg^−1^; P > 0.899). A significant increase occurred during HST2 in HOT (+10 ± 5 mOsm·kg^−1^, P = 0.010), with no changes during HST1. NGAL did not differ at rest (P > 0.05) but increased post-HST only in TEMP during HST1 (+45 ± 29 μg·L^−1^, P = 0.030). KIM-1 showed no within-trial changes in either group during HST1 or HST2 (P > 0.999).

## Discussion

The primary aim of this study was to determine whether five days of isothermic HA could reduce thermal and cardiovascular strain and attenuate renal stress during exercise in the heat. The key finding of this study is that five days isothermic HA improved cardiovascular and thermoregulatory responses, evidenced by reductions in resting and peak heart rate, plasma volume expansion (+7.3%), and increased sweat loss (+500 mL), but did not alter neuroendocrine markers (copeptin, normetanephrine) or renal tubular stress markers (KIM-1, NGAL) during exercise in hot-humid conditions.

Cardiovascular adaptations in the current study are consistent with previous reports of short-term HA benefits but exceeded the mean plasma volume change (+4.3%) reported in earlier studies (Tyler et al., 2016). Cardiovascular improvements likely reflect enhanced stroke volume secondary to plasma volume expansion (Périard et al., 2016), although no change in sweat sodium concentration suggests mechanisms beyond aldosterone-mediated sodium retention, possibly involving isosmotic fluid shifts from the cutaneous vascular space (Senay et al., 1976). Despite these thermoregulatory and cardiovascular benefits, markers of sympathetic activity (normetanephrine) and fluid-regulatory stress (copeptin) remained unchanged. This contrasts with earlier reports of reduced norepinephrine following HA (Hodge et al., 2013; Stacey et al., 2018), suggesting that heart rate reductions may reflect improved cardiac function rather than diminished sympathetic drive (Parsons et al., 2020). Evidence on neuroendocrine adaptation remains equivocal: Stacey, et al., (2018) found no significant change in copeptin after 23 days of acclimatization despite reduced cardiovascular strain, indicating limited AVP-mediated fluid regulation. In contrast, Snape et al., (2025) reported decreases in copeptin and normetanephrine after 7 days of mixed-method HA, highlighting protocol-specific effects.

Improvements in cardiovascular efficiency have often been associated with reductions in core temperature; however, resting Tgi did not change significantly in this study (-0.2 °C), which may reflect high inter-individual variability and independence of different HA indices (Corbett et al., 2018). Therefore, classifying whether a HA intervention has been successful should be done with reference to multiple specific indices of HA, rather than a global classification. Increased sweat loss observed here (+37%) was more pronounced than previous short-term HA studies (+5%) (Tyler et al., 2016), likely due to the high humidity (70% RH) reducing evaporative cooling and promoting greater sweat gland responsiveness (Patterson et al., 2004a).

To our knowledge this is the first study to examine these specific renal and neuroendocrine markers in the context of short-term isothermic HA. NGAL and KIM-1 remained unchanged, indicating that short-term HA does not confer renal protection during exercise in hot-humid conditions. These findings align with Connor et al., (2025) who reported that 10 days of HA improved thermal tolerance but did not attenuate increases in serum creatinine or NGAL during exercise-heat stress, suggesting persistent renal sensitivity to hyperthermia. Furthermore, Ravanelli et al., (2021) observed that seven days of passive HA did not improve glomerular filtration rate during heat stress, although albuminuria incidence was reduced, suggesting better preservation of renal filtration integrity under thermal stress. Pryor et al. (2020) also reported that six day’s HA did not reduce serum creatinine rise, though fewer participants met AKI thresholds post-HA. This aligns with Omassoli et al., (2019) who demonstrated reductions in AKI risk with gradual acclimatisation, and a significant reduction in serum creatinine post-exercise by day 23, reinforcing that renal adaptation is gradual and protocol-dependent.

Collectively, these studies indicate that HA whether short-term, extended, or passive, doesn’t fully protect renal function during heat stress, but may reduce risk of developing AKI. In the current study, it was postulated that reductions in neurohumoral markers (copeptin, NMET) following HA may have helped keep blood vessels in the kidneys more open during exercise-heat stress, improving blood flow and reducing the risk of damage to the kidneys filtering units. However, despite an increase in PV there was no change in copeptin or NMET. This may suggest similar levels of renal perfusion and risk of tubular injury following acclimation. Our findings, highlight that while short-term HA improves cardiovascular and thermoregulatory responses, extended HA protocols (≥10 days), combined with individual hydration plans or cooling interventions may be required to safeguard renal function during exercise in hot environments. This aligns with evidence from occupational settings, were prolonged work in the heat without adequate hydration or cooling increases the risk of kidney injury among workers in physically demanding roles (Schlader et al., 2019). Similarly, athletes undertaking frequent heat training sessions (five times per week for five weeks) for increases in haemoglobin mass and exercise performance (Rønnestad et al., 2021), could experience cumulative renal stress, highlighting the need for structured acclimation and recovery strategies to mitigate potential injury.

This study has several limitations that should be considered when interpreting the findings. Firstly, we didn’t include direct measures of kidney function (e.g. serum creatinine or glomerular filtration rate). While this was intentional to focus on emerging early tubular injury markers (NGAL and KIM-1) and neuroendocrine indices (copeptin, NMET), it limits inferences about acute changes in filtration and the functional significance of any biomarker shifts. Second, kidney injury markers were assessed only in blood. Although circulating concentrations of NGAL and KIM-1 offer practical sampling during repeated exercise-heat exposures, they are not kidney-exclusive and may be influenced by extra renal sources and systemic inflammation. Future research should include a concurrent urinary biomarker panel to better understand tubular site specificity. Finally, while five days HA elicited physiological adaptations in the present study, high inter-individual variability in HST-induced responses combined with modest sample size may have reduced statistical power, increasing the risk of biomarker-specific false negatives. Thus, some “no change observed” outcomes may reflect an underpowered design rather than true absence of effect. This interpretation is supported by prior work (Snape et al., 2025), where even a short HA intervention (+2 days) induced measurable reductions in fluid-regulatory strain. Future studies should consider larger cohorts and/or extended HST protocols to improve sensitivity for detecting renal and neuroendocrine adaptations.

## Conclusion

Five days isothermic HA was effective at evoking meaningful physiological and perceptual adaptations at rest and during exercise-heat stress but did not modify the response of surrogate markers of sympathetic activation, fluid regulation or renal stress. This HA protocol offers a time-efficient and effective strategy for endurance-trained individuals competing in a hot-humid environment. However, longer HA protocols may be required to induce a more complete heat adapted phenotype, which could confer greater protection against renal injury and/or AKI risk during exercise heat stress in athletic and military populations.

## Acknowledgements

The figures were created with GraphPad Prism (version 9) and are published with permission.

## Notes

**Funding** This study was supported by Leeds Beckett University.

### Competing Interest Statement

The authors have declared no competing interest.

